# Macaque Brainnetome Atlas: A Multifaceted Brain Map with Parcellation, Connection, and Histology

**DOI:** 10.1101/2022.10.18.512488

**Authors:** Yuheng Lu, Yue Cui, Long Cao, Luqi Cheng, Zhenwei Dong, Changshuo Wang, Youtong Liu, Baogui Zhang, Haiyan Wang, Kaixin Li, Liang Ma, Weiyang Shi, Wen Li, Yawei Ma, Zongchang Du, Jiaqi Zhang, Hui Xiong, Na Luo, Yanyan Liu, Xiaoxiao Hou, Xinyi Liu, Hongji Sun, Jiaojian Wang, George Paxinos, Zhengyi Yang, Lingzhong Fan, Tianzi Jiang

**Author notes:** Corresponding Authors: Professor Tianzi Jiang, Brainnetome Center, Institute of Automation, Chinese Academy of Sciences, Beijing 100190, China., Professor Lingzhong Fan, Brainnetome Center, Institute of Automation, Chinese Academy of Sciences, Beijing 100190, China., Associate Professor Zhengyi Yang, Brainnetome Center, Institute of Automation, Chinese Academy of Sciences, Beijing 100190, China. The authors contributed equally (Yuheng Lu, Yue Cui, Long Cao). **Authors Contributions** T.J. proposed the concept and designed the protocol. Y.L., Y.C., L.C., performed the experiments and analyzed the data. H.S., Y.L., H.X., and X.L. prepared the ex vivo samples, and B.Z. performed the MRI scanning. C.W and Z.Y. developed the website. G.P., L.F., and Y.L. contributed to the nomenclature. T.J., L.F., and Z.Y. led the project and supervised the experiments. All authors contributed to the writing of the manuscript.

## Abstract

The rhesus macaque (*Macaca mulatta*) is a crucial experimental animal that shares many genetic, brain organizational, and behavioral characteristics with humans. A macaque brain atlas that identifies anatomically and functionally distinct regions is fundamental to biomedical and evolutionary research. However, even though connectivity information is vital for understanding brain functions, a connectivity-based whole-brain atlas of the macaque has not previously been made. In this study, we created a new whole-brain map, the Macaque Brainnetome Atlas (MacBNA), based on the anatomical connectivity profiles provided by high angular and spatial resolution ex vivo diffusion MRI data. The new atlas consists of 248 cortical and 56 subcortical regions as well as their structural and functional connections. The parcellation and the diffusion-based tractography were comprehensively evaluated with multi-contrast MRI, invasive neuronal-tracing, and Nissl-stained images collected from a single subject and with open-access datasets from other cohorts. As a demonstrative application, the structural connectivity divergence between macaque and human brains was mapped using the Brainnetome atlases of those two species to uncover the genetic underpinnings of the evolutionary changes in brain structure. The resulting resource includes (1) the thoroughly delineated Macaque Brainnetome Atlas (MacBNA), (2) regional connectivity profiles, (3) the postmortem high resolution macaque diffusion and T2-weighted MRI dataset (Brainnetome-8), and (4) multi-contrast MRI, block-face, and section images collected from a single macaque. MacBNA can serve as a common reference frame for mapping multifaceted features across modalities and spatial scales and for integrative investigation and characterization of brain organization and function. Therefore, it will enrich the collaborative resource platform for nonhuman primates and facilitate translational and comparative neuroscience research.

## Introduction

Rhesus macaques have been widely employed to explore the mechanisms of human cognitive functions and to model human brain disorders due to their high similarity in genetics, physiology, and brain structure^1, 2^. International brain projects^3–5^ utilize non-human primate research as a core source for elucidating the neural basis of cognition and for promoting translational medicine. A macaque brain atlas that delineates the heterogeneous spatial organizations of the brain can provide a link that can be used to translate findings from rhesus macaques to humans. It can characterize distinct regional profiles, including connectivity, architecture, and topography^6, 7^, and is fundamental to understanding the function, development, and evolution of the brain^8^. The profiles of different brain regions measured by MRI, tracers, and histology uncover the diverse neurobiological properties on different levels^9–11^. So far, only fragmented studies of macaque atlases on a single level, which can reflect only limited aspects of brain organization and function, have been reported. A new macaque brain atlas with a unifying map of parcellations, connections, and histology across scales and with extensive validation has remained elusive.

Macaque brain cartography has as long a history as that of humans^12^, and researchers have created various macaque brain atlases in past decades^13–16^. Previous cyto-, myelo-, and chemoarchitectonic delineations of the macaque brain were primarily developed using neuroanatomical methodologies, and the boundaries of brain structures were determined by microstructural anatomical features^14–19^. Unlike traditional histological methods, animal MRI allows for the intact measurement of the brain. Connectivity-based parcellations have emerged to segregate the brain into distinct regions using diffusion and functional MRI^20, 21^. Since anatomical connections are predictors of functional activations from the primary somatosensory^22, 23^ to the high-order association cortices^24^, identifying the functional diversity of brain regions has been constrained by anatomical connections^25^. Anatomical connectivity-based techniques capture global properties by estimating the physical connections between regions, allowing for a more effective characterization of both the areal and network-level organizations^7^. To the best of our knowledge, a macaque whole-brain atlas based on anatomical connectivity has, until now, been unavailable.

Comprehensive atlases have successfully delineated brain regions based on anatomical connectivity profiles in other primates, specifically humans and marmosets^26, 27^. In rhesus macaques, previous studies using anatomical connectivity to define functional borders mainly focused on particular regions, including the prefrontal cortex^28^, inferior parietal lobule^29^, accumbens nucleus^30^, and other areas^31, 32^. As a result of the limited scan tolerance of macaques in vivo, we collected high quality postmortem diffusion MRI (dMRI) data to provide a high spatial and angular resolution, something that would not have been possible in an in vivo study. The greater amount of time available in postmortem studies allows for a more precise characterization of the connectivity profiles between regions^33, 34^ and, thus, benefits connectivity-based parcellations. Accumulating evidence suggests that integrating anatomical connectivity with multi-modal information such as cytoarchitecture and tracers may be vital for elucidating the specialization of brain regions at multiple levels and scales^10, 11, 35^. The relationship between connection-based and cytoarchitecture-based parcellations has rarely been explored although some studies have implied that they converge in some brain regions^9, 28, 36^. MRI, tracer, and Nissl-stained sections of a single monkey, as utilized in this study, could help to link the macroscopic parcellation and connections with microscopic architectures^37^.

In the current study, we present a fine-grained anatomical connectivity-based macaque brain atlas, the Macaque Brainnetome Atlas (MacBNA), using high spatial and angular resolution dMRI images of 8 postmortem monkeys acquired with an ultra-high field 9.4T MRI scanner. We also present a set of tracer and Nissl-stained sections from another monkey brain to integrate the MacBNA with meso-scale connections and cytoarchitectonic properties. As an application of the uniquely-constructed Brainnetome macaque and human atlases, we mapped the structural divergency map of those two species based on connectivity and studied how the map relates to transcriptomes. We will release the resource, including the MacBNA and the corresponding multi-modal connectomes, the postmortem monkey dMRI data, the block-face images, and the section images, for open access.

## Results

An overview of the data, mapping, and evaluation of the Macaque Brainnetome Atlas is summarized in Fig. 1. First, diffusion MRI data with high angular and spatial resolutions from eight ex vivo monkeys (Brainnetome-8) (Fig. 1A, left panel) were acquired from a 9.4 T MRI scanner. The advantages of Brainnetome-8 were quantitatively compared with two other macaque datasets, Oxford-I^38^ and TVB^39^, from open resources. Tracer and Nissl-stained sections from another monkey, R04, (Fig. 1A, right panel) were acquired for the subsequent evaluation of the parcellation and connections. Second, we constructed a fine-grained connectivity-based atlas of the macaque brain, MacBNA, using the Brainnetome-8 (Fig. 1B, left panel). Then, the structural and functional connectomes were identified based on the MacBNA using Brainnetome-8 and Oxford-II^40^, respectively (Fig. 1B, middle panel). We mapped the cross-species structural divergence based on connectivity using the Brainnetome atlases for macaques (MacBNA) and humans (HumBNA) and explored the genetic mechanism underlying the evolutionary changes in brain connectivity (Fig. 1B, right panel). Finally, we evaluated the MacBNA in three ways: 1) The cyto-boundary of the R04 Nissl-stained images was compared with the MacBNA at a microstructural level (Fig. 1C, left panel). 2) The diffusion-based structural tractography was validated with neuronal-tracing connectivity (Fig. 1C, middle panel). 3) The distance-controlled boundary coefficient (DCBC) was used to compare the MacBNA with other macaque atlases (Fig. 1C, right panel).

**Fig. 1.**
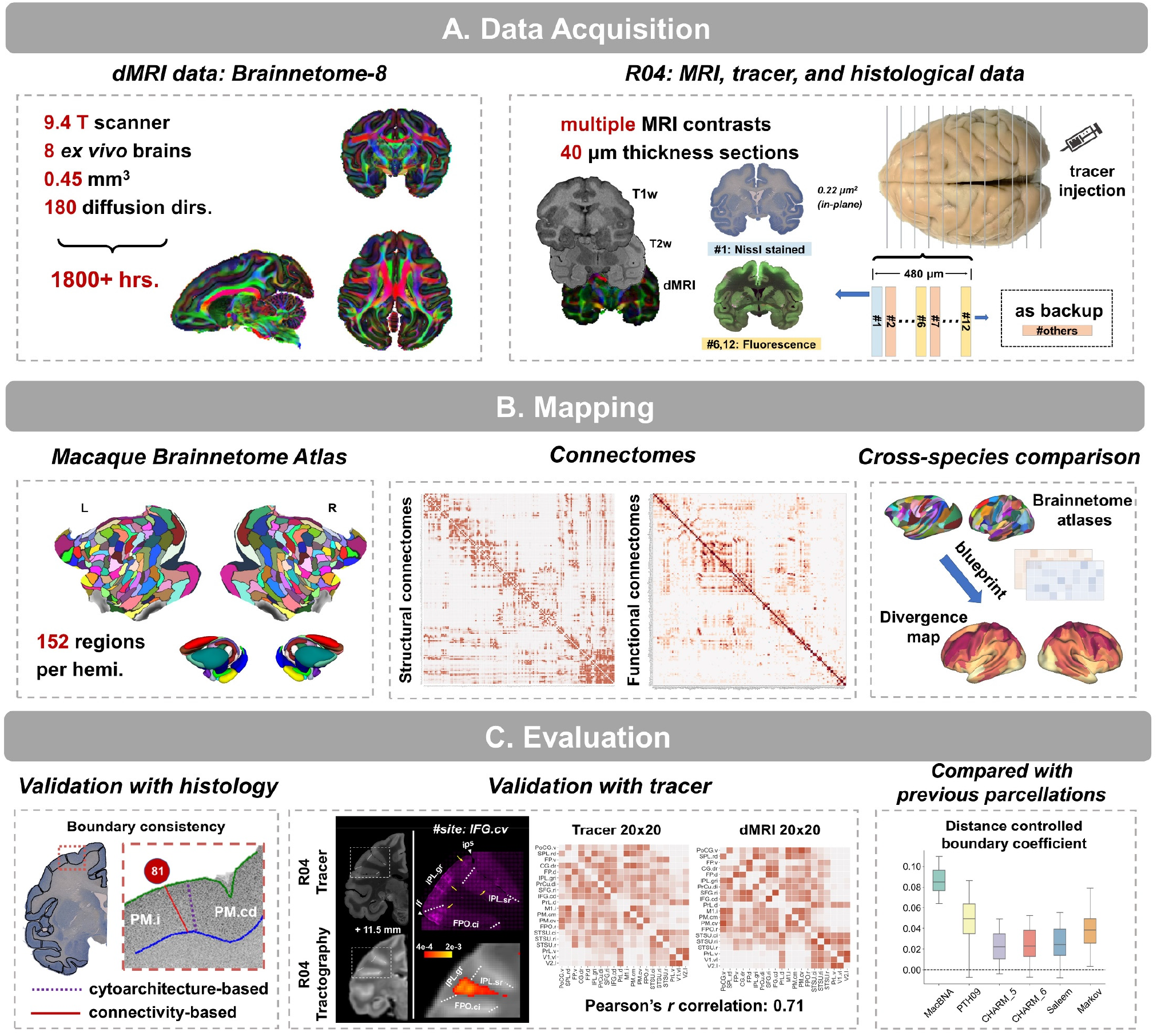
Overview of the data, mapping, and evaluation of the Macaque Brainnetome Atlas. **(A)** The high quality dMRI dataset Brainnetome-8 with eight ex vivo macaque brains and additional collected multi-modal and multi-scale data, including T1w, T2w, and diffusion MRI, Nissl-stained, and fluorescent tracer images for macaque R04. **(B)** A fine-grained structural connectivity-based atlas for macaques, i.e., Macaque Brainnetome Atlas (MacBNA) was delineated using high-quality Brainnetome-8 data, and regional structural and functional connectivity profiles were determined. An evolutional divergence map was constructed with structural connectivity blueprints based on the macaque and human Brainnetome atlases. **(C)** The Macaque Brainnetome Atlas was compared with cytoarchitectonic boundaries using the Nissl sections from R04. The structural connections were evaluated using the tracer images from R04 and an open-access tracer dataset.

### Quality Assessment of Brainnetome-8

High quality dMRI images allow for fine-detailed identification of white matter, leading to accurate estimation of structural connections. Eight ex vivo macaque brains (Brainnetome-8) were soaked with 0.1% gadolinium contrast media for a month to substantially reduce the T1 relaxation time and to improve the quality of the MRI images before scanning with a 9.4 T Bruker MRI scanner. Then, high spatial and angular resolution dMRI data (0.45^3^ mm, 180 multi-shell directions) were acquired for a total of more than 1800 hours. Two other public available macaque datasets, Oxford-I (ex vivo, 0.6^3^ mm, 120 single-shell directions) and TVB (in vivo, 1.0^3^ mm, 64 single-shell directions), were used for comparison with the Brainnetome-8. A gray-matter structure (Fig. 2A) that bridges the caudate nucleus and putamen and can be recognized in the Brainnetome-8 was attributed to the high spatial resolution. The primary orientation of the fibers within the bridge in the Brainnetome-8 revealed that it connected the two subcortical nuclei, but the fibers were either invisible or unclear in the other two datasets. The high resolution could reduce the partial volume effect (PVE) and help dissect tracts from crossing fibers. A tract that was located lateral to the inferior longitudinal fasciculus (ILF) originated from the caudal part of the ILF and then coursed vertically, emerging to the caudal part of the superior longitudinal fasciculus (SLF)^41, 42^. It was visually identified in Brainnetome-8, but it was hard to dissect in either the Oxford I or the TVB (Fig. 2B). More accurate tractography could be ensured because the Brainnetome-8 had the highest ratio of voxels that contained three fibers (i.e., crossing fibers) and the lowest uncertainty of the orientation estimation using FSL (Figs. 2C, D). A 29×91 tracer connection matrix from an open resource^34, 43^ was used to elucidate the differences across the three datasets taking the distances between regions into consideration (bin step = 1 mm). The correlation (Pearson’s *r*) between the dMRI and the tracer connections showed that the Brainnetome-8 improved the identification of long-range connections although its correlation was similar to that of the Oxford-I for distances less than 20 mm. Also, the ex vivo datasets had substantial advantages over the in vivo one at all distances (Figs. 2E, F).

**Fig. 2.**
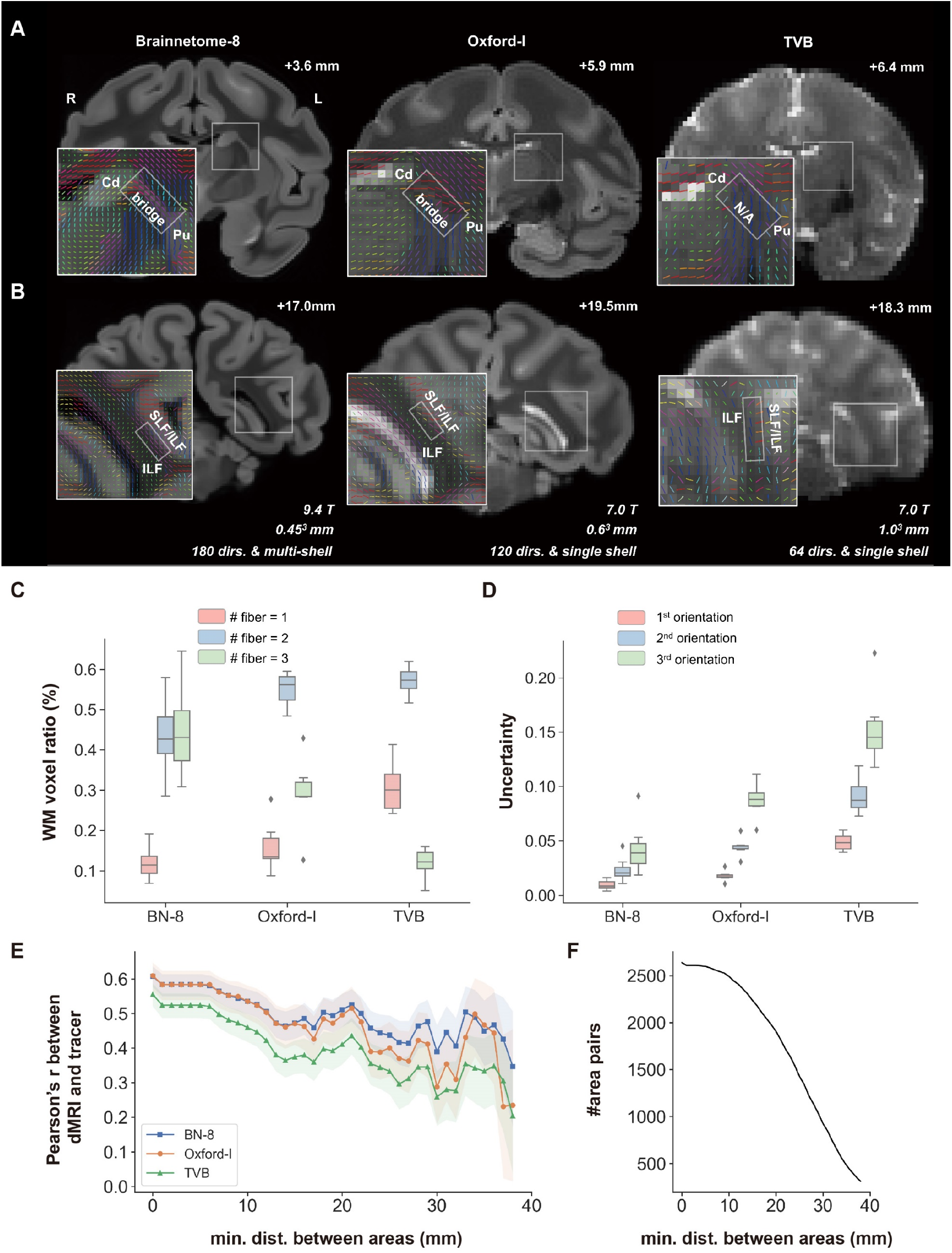
White matter structural comparisons of the Brainnetome-8, Oxford-I, and TVB dMRI datasets. **(A-B)** Visual inspection of the three datasets shows a substantial improvement of the Brainnetom-8 over the other datasets in that it captured the brain structures better. This can be attributed to its high spatial and angular resolution compared with the Oxford-I and TVB datasets. The structure highlighted in the white box is zoomed in at the left bottom corner of each coronal slice. **(C)** The ratio of white matter voxels that contain 1/2/3 fibers and **(D)** the uncertainty of each estimated fiber orientation provided by Bedpostx, indicating that more crossing fibers could be resolved and a sharper distribution of fiber orientations was estimated with Brainnetome-8 **(E)** A higher correlation was found between the connections with Brainnetome-8 dMRI and tracer, taking the minimal distance between regions into consideration. Solid lines and shadings denote mean and standard deviation of the bootstrapped distributions, respectively. **(F)** The number of all paired regions at varying minimum interareal distance is shown. Abbreviations: *BN-8, Brainnetome-8; Cd, caudate nucleus; Pu, putamen; SLF, superior longitudinal fasciculus; ILF, inferior longitudinal fasciculus*.

### Structural Connectivity-based Parcellation of Macaque Brains

Using the 8 ex vivo high-quality dMRI dataset, we created a fine-grained whole-brain parcellation of macaque brains based on structural connectivity. Following the protocols used in the HumBNA construction^26^, the whole brain was first delineated into 29 initial seed units, each of which was parcellated into multiple subdivisions based on its connectivity with the optimal cluster number determined by three cluster indices (Figs. 3A-F). The whole brain was finally divided into 248 cortical and 56 subcortical subregions with each hemisphere having an equal number of regions (Figs. 4A, B). To evaluate the validity of the MacBNA, DCBC was used to measure the homogeneity and heterogeneity of the parcellations using structural connectivity. DCBC is an unbiased criterion that can compare within-parcel to between-parcel similarity considering the spatial autocorrelation of the brain data and the scale of the parcellations. In other words, a higher DCBC indicates that the parcellation provides a brain topography with higher within-parcel similarity and higher between-parcel dissimilarity. We compared the DCBC of the MacBNA using the structural connectivity of each vertex/voxel with other existing parcellations of macaques, including Paxinos et al.’s^18^, CHARM^19^, Saleem et al.’s^44^, and Markov et al.’s^43^. The results in Fig. S8 demonstrate that MacBNA achieved the best performance for the delineation of the macaque brain based on structural connectivity (paired *t* test, FDR corrected *P* < .0001). We also examined whether there is bias in the distribution of the parcellation boundary located in the gyri or sulci when using the ‘sulc’ map generated by Freesurfer^45^. The ratio of the MacBNA boundary vertices that are located in gyral regions (sulc > 0) to those in sulcal regions (sulc < 0) was not significantly greater than random using 10000 permutations (*P*_spin_ > .05) (Fig. S16).

**Fig. 3.**
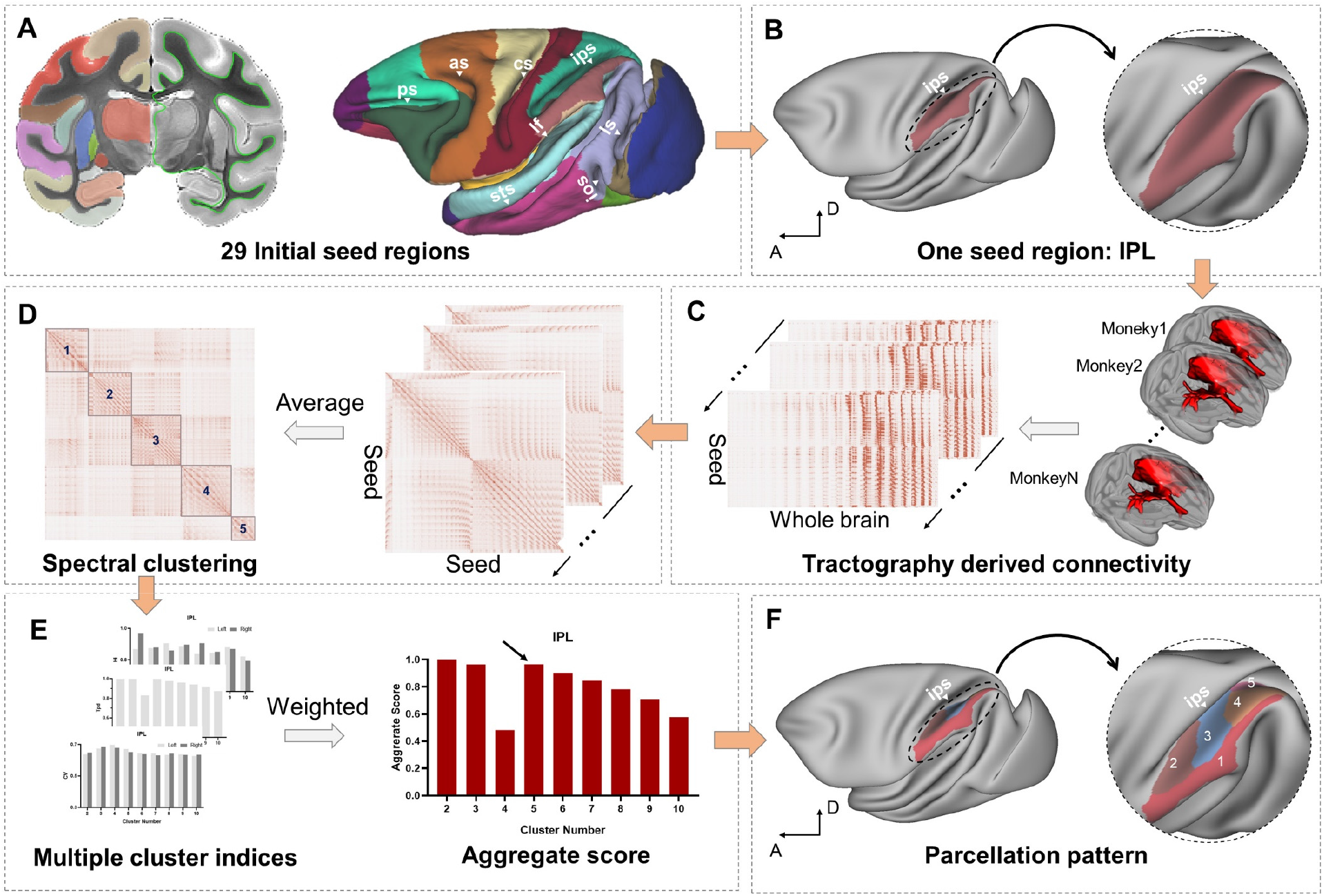
Pipeline for the construction of the Macaque Brainnetome Atlas. **(A)** 29 initial seed regions of interest (ROIs), including 21 cortical and 8 subcortical regions, were manually delineated in volume space. The coronal view of the template brain shows the location of the anatomical seed regions as colored areas (left hemisphere) and as green lines on the white/gray surface (right hemisphere). Those seed regions were then mapped to the surface space to get surface-based ROIs for the cortical parcellation. (**B-F)** Surface-based parcellation of an exemplar seed region, IPL. **(C)** Probabilistic tractography was performed from each vertex seed to the whole brain and the connectivity profile was stored in a connectivity matrix (seed-by-whole brain) for each subject. **(D)** Pairwise correlations of voxels in the seed region were calculated to get a similarity matrix (seed-by-seed) for each subject, and spectral clustering was conducted on the averaged similarity matrix across all subjects. **(E)** Three indices, CV, Tpd, and Hi, were fused to form an aggregate score based on entropy theory to estimate the optimal cluster number (5 was the optimal cluster number as indicated by the location of the local maximum). **(F)** The parcellation pattern of the IPL. Abbreviations: *as, arcuate sulcus; cs, central sulcus; ps, principal sulcus; lf, lateral fissure; sts, superior temporal sulcus; ls, lunate sulcus; cgs, cingulate sulcus; ots, occipitotemporal sulcus; pos, parieto-occipital sulcus; iocs*, *inferior occipital sulcus*; *IPL, inferior parietal lobule*.

**Fig. 4.**
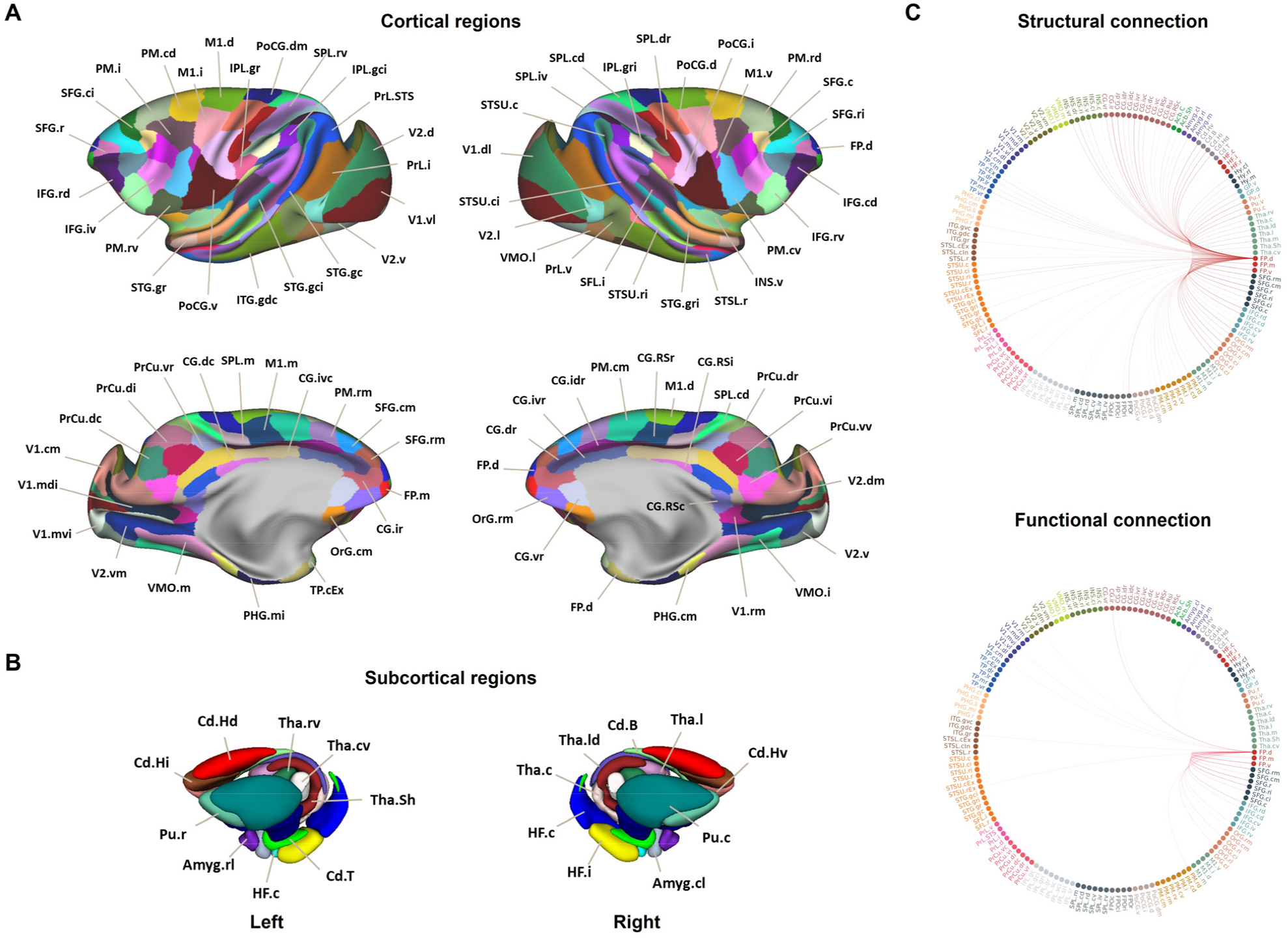
Overview of the Macaque Brainnetome Atlas. **(A)** Surface view of 248 cortical regions and **(B)** 3D view of 56 subcortical regions. **(C)** The left intra-hemisphere structural and resting-state functional connection of an exemplar region, FP.d. The nomenclature information for each subregion is listed in the Supplemental Nomenclature. Abbreviation: *FP.d, frontal pole dorsal part*.

### Multifaced Mappings and Evaluation of Premotor Cortex

Taking the premotor cortex (PM) as a representative instance (another example of the OrG in Fig. S4, S5), we identified seven subregions, the PM.rd, PM.cd, PM.i, PM.cv, PM.rv, PM.rm, and PM.cm, based on the local maximal positions of their clustering indices (Fig. 5A, left panel) (Fig. S2A). Using the data from the multiple subjects collected in Brainnetome-8, we calculated the population probabilistic map of each subregion to reveal the individual variation (Fig. 5A, middle panel), which indicates the stability of the parcellation map across subjects. The structural connectivities of the PM.cd and PM.i (Fig. 5A, right panel) show the differences in their connection patterns with the whole brain. Specifically, the PM.cd showed stronger connections with the medial cortex and posterior intraparietal regions, and the PM.i had stronger connections with the ventral premotor and sensorimotor areas.

**Fig. 5.**
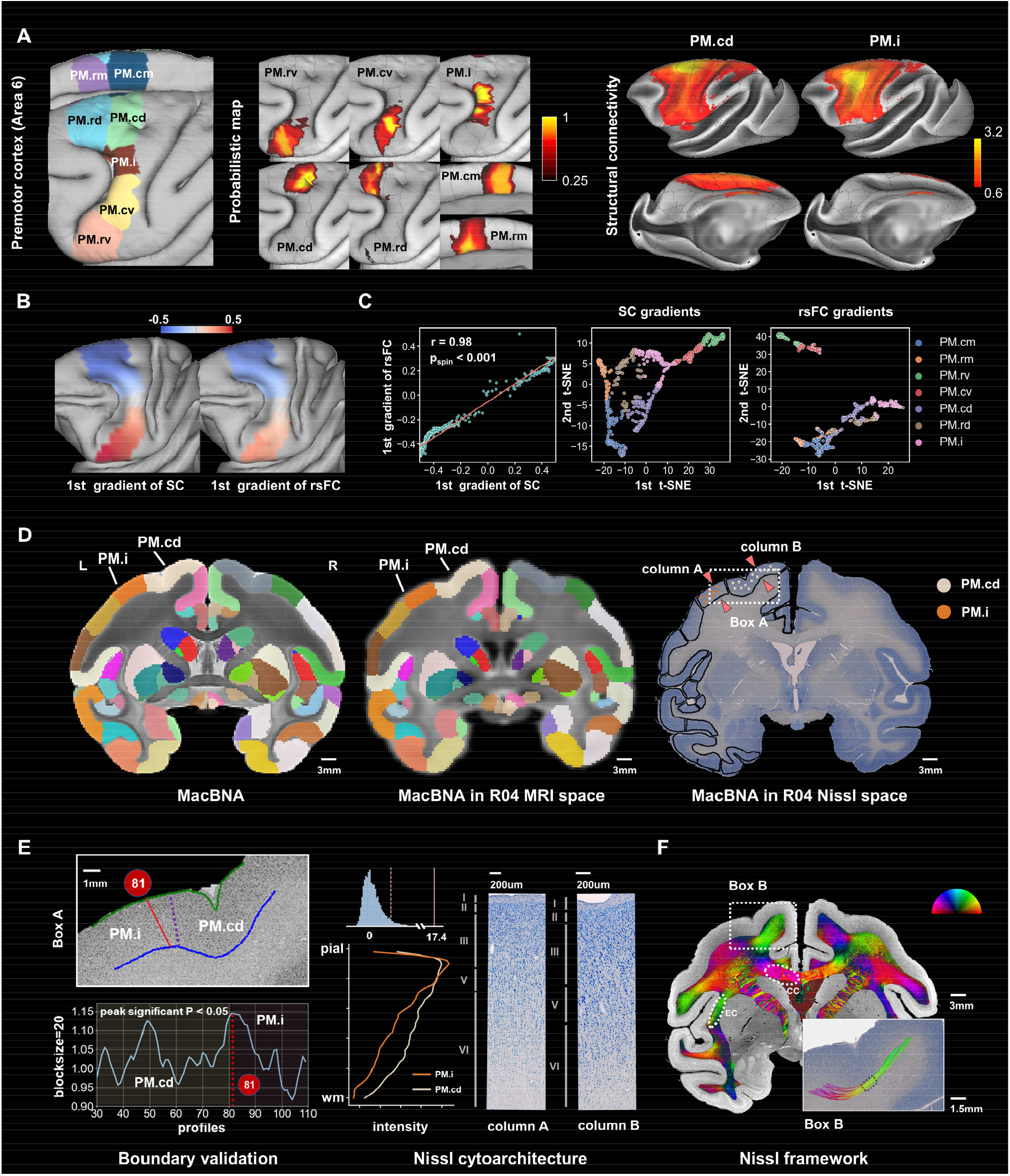
Macaque Brainnetome Atlas of the premotor cortex (PM) and the exemplar joint study with the MRI and Nissl images. (A) Left panel shows the parcellation of the PM in the MacBNA, which subdivided it into 7 regions. Middle panel is the population probabilistic map of each subregion of the PM, indicating the individual variation. Right panel is the structural connectivity of the PM.cd and PM.i with the whole brain. **(B)** The first gradient of the structural and resting-state functional connectivity of the PM. **(C)** The correlation between the two gradients and the visualization of the 7 clusters of the PM with the t-SNE. **(D)** Parcellations of the MacBNA in template, R04 MRI, and R04 Nissl space. The gray and orange dots mark the regions of the PM.cd and PM.i, respectively. **(E)** The zoom-in details (marked with white box) indicate the boundary between the PM.cd and PM.i of the MacBNA (purple line) based on connectivity and the boundary (red line) determined using a quantitative cytoarchitecture analysis (left bottom). The middle panel is the mean density profile of the PM.cd and PM.i regions, and a permutation test with 10000 samples indicates the significant difference between their profiles (*P* < .001). The right panel shows the detailed laminar structures of two selected columns of the two regions that were marked in (D). **(F)** The fiber orientation reconstructed with the glial framework at a mesoscale spatial resolution (44 microns), and the white circles indicate two major tracts, i.e., CC and EC. The right bottom shows a tiny U-fiber connecting the neighboring gyral regions when seeding from the region below the sulci (black dotted box). Abbreviations: *PM, premotor cortex; PM.rd, premotor cortex rostrodorsal part; PM.cd, premotor cortex caudodorsal part; PM.i, premotor cortex intermediate part; PM.cv, premotor cortex caudoventral part; PM.rv, premotor cortex rostroventral part; PM.rm, premotor cortex rostromedial part; PM.cm, premotor cortex caudomedial part. CC,* c*orpus callosum; EC, external capsules*.

A gradient analysis using a diffusion map algorithm^46^ based on the structural and functional connectivity of this premotor region shows that the first gradients of the two modalities had a high correlation coefficient (*r* = 0.98, *P*_spin_ < .001, Figs. 5B, C). For the second and the third gradients, the correlations decreased to *r* = 0.689 (*P*_spin_ < .001) and *r* = 0.205 (*P*_spin_ < .001), respectively (Figs. S2C, D). A 2D visualization of the gradients using t-SNE^47^ for each subregion presented a clustering pattern for both the structural and functional modalities, visualizing our parcellations from the viewpoint of gradients.

To explore the cytoarchitectonic variations between the regions, we collected Nissl-stained sections from a postmortem monkey (ID: R04) and mapped the MacBNA parcellation to the corresponding Nissl slice space (Fig. 5D) with the black boundary delineating the subregions of the MacBNA in the left hemisphere and the white and orange dots marking the respective locations of the PM.cd and PM.i. The density profiles of those two regions (marked as box A in Figs. 5D, E) along the pial and white surfaces was extracted to obtain the cytoarchitecture-based boundary. A sliding window procedure was used to calculate the distance between the neighboring blocks (see **Methods** for details). The distance function of the profile positions located the maximal point as a peak with the block size set at 20. The block size was set from 11 to 24 to ensure that the maxima were robust and independent of this parameter (Fig. S3A). A Hotelling’s T^2^ test was used to confirm the significance of the peak (*P* < .05), and the two neighboring sections underwent the same procedure to ensure that the peak location was stable and not a false positive caused by artifacts or outliers (Figs. S3B, C). The cyto boundary based on the profile density analysis (position 81, red line in Fig. 5E left panel) was close to the boundary determined by the connectivity (purple line) (distance between boundary centers = 0.79 mm). Fig. 5E (middle panel) shows the mean density profiles of the PM.cd and PM.i from the pial to the gray/white surface. A permutation test (10000 samples) was utilized to test the significance of the difference in density profiles between the two subregions (*P* < .001, Fig. 7E). A detailed laminar structure of a column of each subregion (Fig. 5E right panel) shows that column A, which was selected from the PM.i (as marked with arrows in Fig. 5D), had a denser layer III while column B, which was selected from the PM.cd, had larger pyramidal cells in layer III. The different aspects of the cytoarchitectonic analysis collectively explained our connectivity-based parcellation at a microscopic level.

The glial framework allows for an estimation of the orientation of the fibers in the white matter at a high spatial resolution (44 microns in our study) and could be used for tracking tiny tracts like U-fibers that might not be reconstructed directly via MRI due to its insufficiently high resolution. Fig. 5F shows the principal orientation organization that was encoded by the RGB color of the stained section using the glial framework, which delineated the anatomical structures of major tracts such as the corpus callosum (CC) along the left-right axis and the external capsule (EC) along the inferior-superior axis (Fig. 5F). A qualitative comparison of the fitting orientation between the DTI and the glial framework revealed their consistency in some major tracts (Fig. S7). A tiny tract was reconstructed directly via the glial framework that connected the neighboring gyral regions when seeded from the area below the sulci (marked in box B in Fig. 5F). Constructing the glial framework takes advantage of the high resolution of the Nissl sections and is useful as a complement to dMRI.

### Comparison of Diffusion-based Tractography and Neuronal-tracing Connectivity

Three injection sites, the IFG.cv, SFG.ri, and IFG.cd, of R04 were coordinated with the MacBNA as the reference system (Fig. 6A). We found that one of those three targets (IFG.cv) had long-range tracer connections with the inferior parietal regions, such as IPL.gr, IPL.sr, and FPO.ci, while the other two did not. A seed ROI was identified as the intersection of a sphere (radius = 3 mm) centered at the coordinates of the injection site with the white/gray surface in the R04 native diffusion space for subsequent probabilistic tracking. The trajectory of the tractography indicates that more streamlines from the IFG.cv terminated around the IPL regions, consistent with the tracer data (Fig. 6B). The contrast between the normalized connection strength of the regions identified by the R04 dMRI, as well as that of the connections from the Brainnetome-8, was similar to that of the number of tracer-labeled neurons (Fig. 6C).

**Fig. 6.**
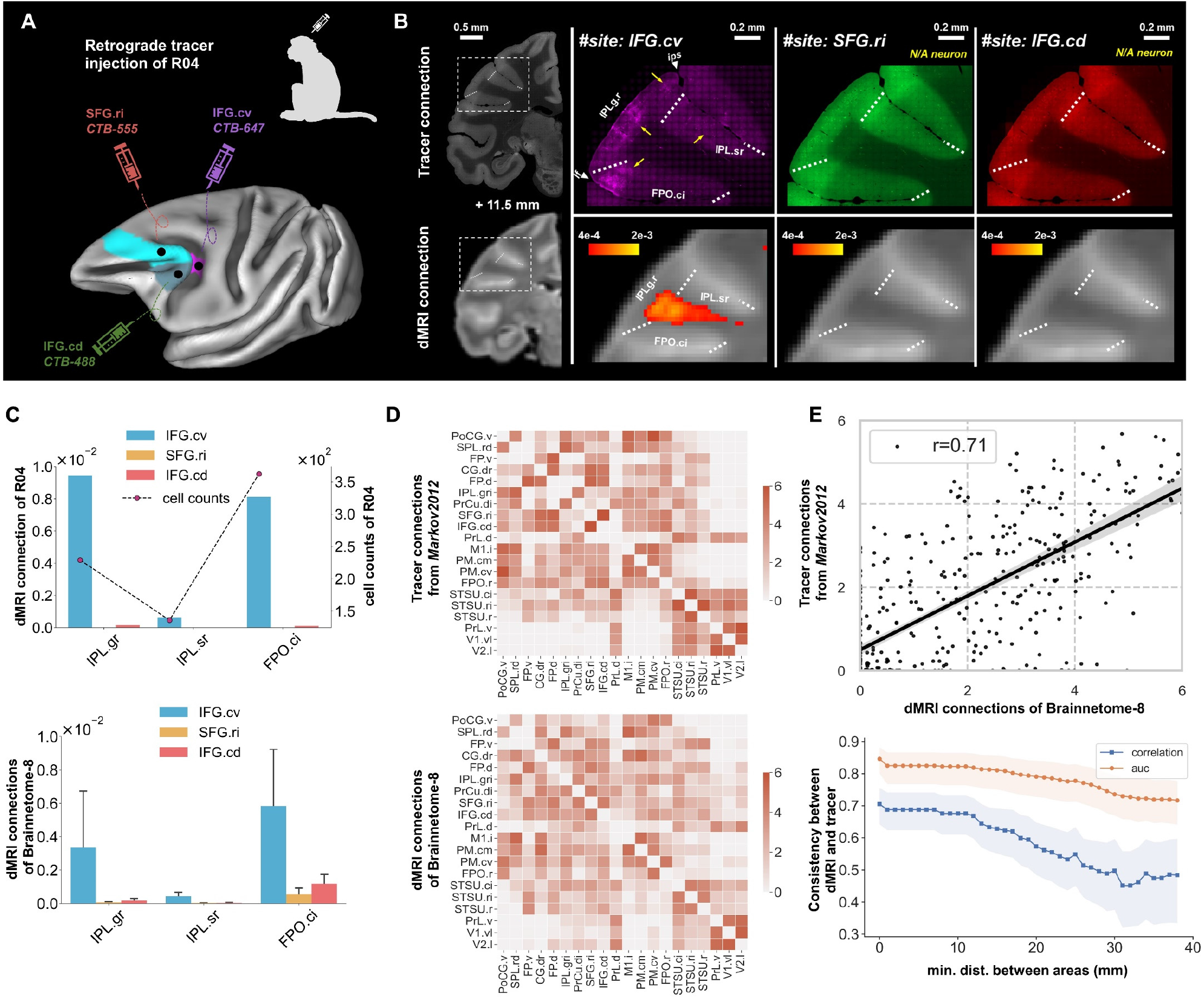
Evaluation of the MacBNA connections with tracers. **(A)** The R04 monkey was injected with retrograde tracers targeted at three regions of the MacBNA, the IFG.cv, SFG.ri, and IFG.cd, indicated with black circles. **(B)** The tracer connections with three projection regions IPL.gr, IPL.sr, and FPO.ci were qualitatively consistent with the tractography results from R04. The labeled neurons were marked with yellow arrows. Then, **(C)** the number of labeled neurons, which connected with IFG.cv, was quantitively compared with the normalized connections based on the dMRI results from R04 and Brainnetome-8. In addition, we compared the dMRI connections based on the MacBNA with the open-access tracer data taking the interareal distances into account. The 20x20 matrices of both the dMRI and tracer show **(D)** a similar pattern and **(E)** a high correlation (*r* = 0.71). Both matrices were also binarized to evaluate the sensitivity and specificity of the dMRI connections. Abbreviations: *IFG.cd, inferior frontal gyrus caudodorsal part; SFG.ri, superior frontal gyrus rostrointermediate part; IFG.cv, inferior frontal gyrus caudoventral part; IPL.gr, inferior parietal lobule gyral rostral part; IPL.sr, inferior parietal lobe, sulcal rostral part; FPO.ci, frontal parietal operculum caudointermediate part; auc, area-under-curve score*.

To reveal the level of agreement between the structural connections and those derived from tracer data more comprehensively, an approximate comparison was made based on the forementioned open-access tracer data, which contains more connections across the whole macaque brain. The injection sites and tracer matrix were mapped to our atlas since the original targets were identified in Markov et al.’s atlas^43^, which has 91 areas. The structural connection based on tractography with the Brainnetome-8 dMRI data was fractionally scaled and log-normalized as in Markov et al.’s study. The connection matrix of the 20 selected regions indicates that the dMRI and tracer data had similar patterns (Fig. 6D), as indicated by a high Pearson’s correlation coefficient of *r* = 0.71, *P* < 10^-6^ (Fig. 6E). The correlation decreased to 0.5 when the areal distance was greater than 20 mm, while the area-under-curve scores between the binarized connections matrices were always above 0.7 when the areal distances ranged from 0 to 40 mm (Fig. 6E). These findings reveal that dMRI was better at identifying short-range connections than long-range connections and had a good sensitivity and specificity regarding whether a connection was present or absent between regions for not only short but also long-range connections.

### Cross-species Structural Connectivity Divergence Based on the Macaque and Human Brainnetome Atlases

The well-delineated parcellations based on MacBNA and HumBNA, which were both based on connectivity, provided a basis for exploring the evolutionary structural connectivity divergence between macaque and human brains. The regional structural connectivities of those two species were represented as the connection probability based on 42 homologous major tracts, i.e., connectivity blueprint^48^ (Fig. 7A). We denoted the structural divergence as the symmetric Kullback-Leibler (KL) distance between the group-wise connectivity blueprints of each region from the MacBNA and each region from the HumBNA. The divergence map represents the minimum divergence of each region of the HumBNA from each region of the MacBNA, indicating that the dorsal prefrontal, posterior parietal, and medial prefrontal cortices had greater divergence, while the visual and temporal cortex had less divergence (Fig. 7B). The divergence map was also significantly correlated with some existing brain maps such as the developmental expansion, cerebral blood flow, and glucose metabolism maps (Fig. 7C) from neuromaps^49^. Furthermore, partial least squares regression (PLSR) was used to investigate the association between the divergence map and the gene expressions using the Allen Human Brain Atlas (AHBH) transcriptome dataset^50^. The first component of the PLSR (PLS1 score, explained variance = 20.4%, *P*_spin_ < .05) had the highest Pearson’s correlation with a divergence map of *r* = 0.47 (*P*_spin_ < .05) (Fig. 7D). The 2257 genes with the most positive and negative weights (|Z| > 3) were enriched in the intelligence, cognition, and mental disease-related gene sets, which had been obtained from DisGeNet^51^ (Fig. 7F). Finally, these 2257 genes were found to be more significantly overlapped (78 genes) with the human-accelerated genes (391 HAR genes remaining in the 15633 AHBH genes set)^52^ using Fisher’s exact test (*P* < .0001) (Fig. 7G).

**Fig. 7.**
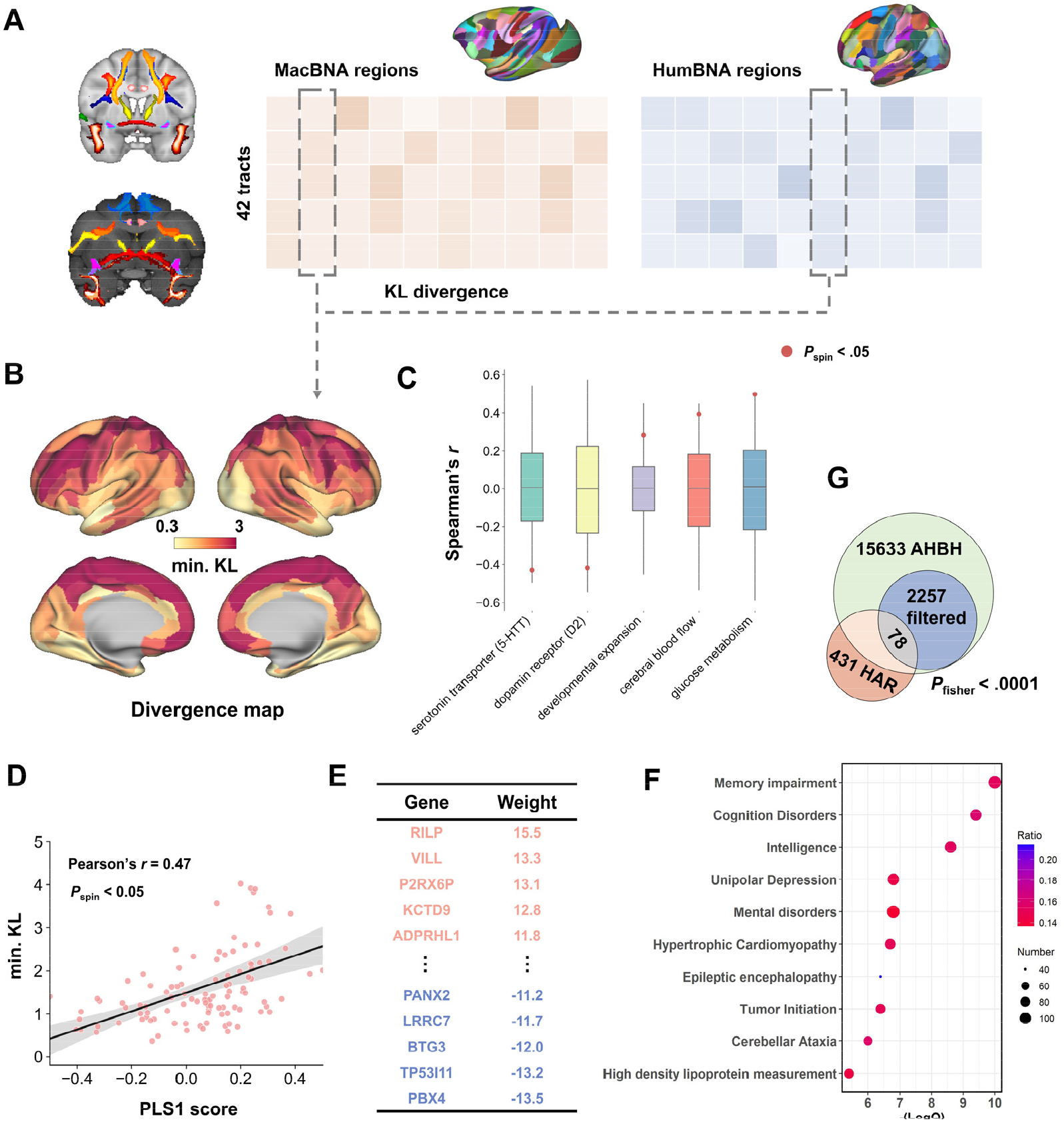
Structural divergence map between macaques and humans with the Brainnetome atlases and its association with transcriptome. **(A)** 42 homologous tracts of macaques and humans were identified to construct a common comparative space to measure the KL divergence between each pair of brain regions of macaques and humans using the blueprint connectivity. (B) The divergence map represents the minimal KL divergence of each region in humans from the macaque regions. (C) We compared the divergence map with other brain maps with neuromaps tool. The points represent the empirical Spearman’s correlation between the divergence map and the other maps (red color indicate *P*_spin_ < .05). The boxplot represents the null distribution of the correlation between the ‘rotated’ divergence map with the other maps (*N* = 10000). (D) Correlation map of the divergence map with the PLS1 score of genes with the AHBH transcriptome dataset (left brain). (E) 2257 genes that have the most positive or negative weights (|Z| > 3) in PLSR were filtered for the subsequent gene ontology analysis. (F) The filtered 2257 genes were enriched in multiple gene sets that are associated with intelligence, cognition, and neuropsychiatric disorders. (G) 78 out of the 431 HAR genes (393 overlapped with 15633 AHBH genes) hwere included in the identified 2257 genes (Fisher’s exact test *P <* .0001)

## Discussion

We produced a 152-area (124 cortical and 28 subcortical areas) per hemisphere macaque brain atlas, MacBNA, based on structural connectivity derived from high-quality ex vivo dMRI data and determined connectivity profiles for each subregion. Conjunct analyses with multi-contrast MRI, neuronal tracing, and cytoarchitectonic data from a single macaque collected in the present study and other independent datasets provided comprehensive and enhanced validations for the brain parcellation and connections. Comparative analyses based on the uniformly well-delineated Brainnetome atlases for macaques (MacBNA) and humans (HumBNA) unveiled the evolutionary hierarchical adaptations in structural connectivity and its genetic underpinnings. In addition, the new MacBNA atlas provides a common coordinate framework for spatially integrating large amounts of data across modalities and scales, providing multifaced maps at various levels including macroscopic brain morphology, structural and functional connectivity, mesoscopic tracer-based connectivity, anatomical microstructures, etc. Therefore, MacBNA, together with the comprehensive data, can be a valuable resource for joint investigations of brain organization and function, which can promote progress in comparative neuroscience and translational medicine.

Brain topography can be characterized from multiple facets including connectivity, cytoarchitecture, and function. Anatomical connectivity is one of the domains that can be used to identify brain spatial heterogeneity, and our atlas preserves the brain topography from the perspective of anatomical connectivity substantially better than other macaque brain parcellations (Fig. S8). Non-human primate studies revealed that cytoarchitecture is partially determined by cortical connectivity^53, 54^ and that the boundaries identified by the anatomical connections between brain areas conform to some degree to the histological subdivisions of the cortex^8, 30, 36^. The subdivisions of the IPL gyral area, as well as those of the dorsal and medial areas of the premotor cortex, in the MacBNA are consistent with existing cyto or myeloarchitectonic atlases^18, 55, 56^. Compared with histological architecture, which characterizes local properties of cell morphology and laminar structure, anatomical connectivity subdivides brain regions by taking advantage of the extensive interconnections within the whole brain that govern the nature and flow of neural activity, reflecting the underlying functional topography of the brain. For example, we segregated the somatosensory cortex into four subregions that appear to represent the face, forelimb, trunk, and hindlimb fields found in a functional somatotopy study^57^ rather than following the cytoarchitectonic parcellations along the rostral-caudal axis (areas 3a, 3b, 1, 2). Area M1 is traditionally considered to be homogeneous within the regions indicated by traditional histological studies^12, 14^, but our parcellation results for the M1 show a pattern along the ventral-dorsal axis, which closely mirrors the somatotopic organization of this area^58, 59^, suggesting that the functional specialization of regions underlies the connectivity^60^.

Since it is widely accepted that there is no golden standard for parcellating the brain, delineating the brain on multiple scales^7, 61^ is essential if we are to explore the brain organization comprehensively. Hence, we provided the MacBNA as a reference system to link the macroscopic connectivity with the microscopic neuroanatomy. By using multi-modal and multi-scale data, various profiles for each region, i.e., the structural and functional connectivity, gradients, and cytoarchitecture, could be mapped (Figs. 5). The connection profiles of each subregion enable the functional role of regions to be tentatively inferred and understood^62^. The heterogeneous connectivity of the regions lying in the peri-arcuate area (Figs. 6B, C) could unveil the functional specificity of those regions. The IFG.cv, which is located in the transition zone between the agranular premotor area 6 and granular area 8, may play a role in controlling the oculomotor as it has a strong connection with the anterior inferior parietal cortex^63^, whereas the granular regions SFG.ri and IFG.cd have only no or weak connection. These latter two regions, the SFG.ri and IFG.cd, which lie anterior to the frontal eye field, may be more related to the top-down control of visual attention^64^. The cytoarchitectonic differentiation between regions could be quantified using the histological data^10^, giving rise to an observer-independent cytoarchitectonic subdivision (Fig. 5E) that could be compared with other parcellations. Observer-independent cytoarchitectonic analysis avoids the traditional visual inspection of stained sections, and researchers could take advantage of the high-resolution histological images to identify finer-grained subdivisions^55, 65^. Since gradient is a popular way to represent brain regions as continuous maps rather than as hard-edged parcellations^66–68^, it can provide an alternative perspective that can help to reflect the neuroanatomy and explain the parcellation pattern in the gradient space (Figs. 5B, C). A recent study indicated that the dorsal premotor cortex has a cognitive-motor behavioral gradient^69^ along the rostral-caudal axis in humans that is consistent with the third structural gradient we found in the PM (Fig. S2). Kahnt et al. proposed that the medial-lateral and posterior–anterior distinctions of the orbitofrontal cortex reflect abstract reinforcers^70^, which is similar to the first and third structural gradient of the OrG that we found (Figs. S4, 5). All of these profiles of the brain are complementary to each other and contribute jointly to the formation of brain function. Focusing solely on one will result in a one-sided view of understanding the structure-function tethering of the brain.

Conducting cross-species comparison in a common space^71^ is commonly done, but it is also crucial to compare the brain organization with parcellations based on unique architecture. The MacBNA and HumBNA, which were uniformly delineated using structural connectivity, provide an alternative to the traditional cytoarchitecture atlases of macaques and humans for uncovering evolutionary changes. The identified divergence map formed using MacBNA and HumBNA revealed that the greatest evolutionary variation in connectivity between humans and macaques is located in the prefrontal and parietal cortices, which are closely related to high-level cognitive functions^72, 73^ and are supported by a higher metabolism rate^74^. However, the fact that the relatively new brain organization of human from an evolutionary timescale perspective may make itself more vulnerable to stress, trauma, or neurodevelopmental conditions, resulting in the psychiatric disorders^75, 76^ that we found genetic evidence for using the AHBH transcriptomes datasets (Fig. 7F). The association between the divergence map and gene expression revealed in this study adds to our understanding of how transcriptional specialization shaped the brain structure, leading to not only higher cognition and intelligence but also to mental related diseases^77, 78^. The accumulation of postmortem dMRI data from primates makes it possible to generalize the parcellation framework to construct connectivity-based atlases^79^, facilitating a polygenetic comparison across primates.

The datasets in this study include not only a new macaque atlas but also its corresponding structural and functional connectomes, as well as high-quality dMRI and tracer data and Nissl-stained sections, making MacBNA a multi-scale multi-modal atlas resource. One of the challenges associated with dMRI is its inability to produce images with high resolution. This low resolution limits the dissection of the white matter bundles and the mapping of the tract pathways in living humans or other primates. However, ex vivo dMRI has the advantage of higher field strength and longer scan duration with tailored pulse sequences^80^ and contrast agents to improve the data quality^27^. Compared with the other two public datasets, Brainnetome-8 could be used to depict the detailed neuroanatomy of not only the major tracts but also tiny/thin tracts because of its relatively high spatial resolution^81–83^. Using Brainnetome-8 enabled us to resolve more crossing fibers within the white matter and sharpen the probabilistic estimation of the fiber orientation (Fig. 2) since using multiple angular contrast and a multi-shell acquisition strategy with high b-values as we did can provide better estimates of different diffusion compartments and lower the uncertainty of the probabilistic fitting of the diffusion orientation^84, 85^. The connections obtained using tracers, tractography, and the glial framework^8–10^ allow for cross-validation by independent aspects of brain properties. Brainnetome-8 performed better than the other released datasets of the ex vivo macaque brain when identifying long-range connections. This is consistent with findings that a higher spatial and angular resolution is important for improving the level of agreement between tracers and dMRI^33^. A recently developed technique^37^ utilized the local organization of glia cells from Nissl stained sections to reconstruct the fiber orientations at a microscale resolution. A qualitative comparison revealed its good correspondence with the dMRI across species^37, 86^, as we found in a single macaque animal (Fig. S7). The extremely high in-plane spatial resolution provided by the glia framework also allowed for the reconstruction of tiny U-fibers (Figs. 5F, S4F), compensating for the lack of resolution in dMRI.

In summary, we present a new macaque brain atlas, the MacBNA, based on structural connectivity with high spatial and angular resolution dMRI images. The MacBNA consists of not only a plausible and fine-grained parcellation but also the detailed macroscopic connections of each subdivision. It can provide a reliable reference system for coordinating and integrating a variety of types of brain maps, including imaging (e.g., multi-contrast MRI, tracing and Nissl-stained data) and genomics (e.g., transcriptomes and proteomes) data to develop a multifaceted brain map of macaque. The multi-modal multi-scale dataset in this study will also provide an open-access platform for addressing computational issues such as modeling a virtual brain and cross scale image registration. We are still in the process of collecting data to further enrich the MacBNA with additional tracer and staining images. In addition, the conceptual framework underlying the construction of the Brainnetome Atlas can be extended to other species for comparative research. Thus, we envision that the Macaque Brainnetome Atlas and the associated dynamic multi-modal multi-scale resources will play a role in cross-species comparisons, translational medicine, and computational modeling.

## Methods

### Animals and Data Acquisition

Eight post-mortem monkeys (Brainnetome-8) were utilized to delineate the atlas and construct the structural connectomes, and another two open access cohorts of 6 ex vivo animals (Oxford-I) and 8 in vivo animals (TVB) were used for validation. Another open access in vivo dataset (Oxford-II) was used to map the resting-state functional connectomes. An additional female adult macaque was scanned to acquire multi-contrast MRI, neuronal tracing, and Nissl-stained histological images covering the whole-brain. All animal procedures in this study were approved by the Ethics Committee of the Biomedical Research of Institute of Automation, Chinese Academy of Sciences (No. IA-202032).

#### MRI of Brainnetome-8 and Open-access Datasets

The Brainnetome-8 consisted of 2 male and 6 female (age = 5.6±1.06 years old) adult monkeys (*Macaca mulatta*). Each monkey was intraperitoneally injected with an overdose of pentobarbital. After a careful confirmation of deep anesthesia, a transcardial perfusion was performed with 500 ml of phosphate-buffered saline (PBS) solution containing 1% heparin (pH 7.4) and 0.1% gadolinium-diethylenetriamine pentaacetic acid (Gd-DTPA). After this 500 ml perfusion, the body organs such as the heart and bladder were removed, and the perfusion rate was decreased to 30 ml/min. This was followed by perfusing with a PBS solution containing 4% paraformaldehyde and 0.1% Gd-DTPA. After two hours of perfusion, the head was removed from the body, and the skull was carefully stripped to expose the whole brain. The formalin-fixed macaque brain was soaked in 0.1% Gd-DTPA and 0.1% sodium azide under sterile conditions for 4 weeks to reduce the T1 relaxation time. Prior to imaging, the specimens were transferred to an MRI compatible holder bundled with medical gauze and immersed in Fomblin (Solvay, Brussels, Belgium) to prevent dehydration and susceptibility artifacts; air bubbles on the brain surface were removed with an aspirator.

Ex vivo MRI data were acquired on an ultra-high field 9.4T horizontal animal scanner (Bruker Biospec 94/30 USR, Ettlingen, Baden-Württemberg). The gradients were equipped with a slew rate of 4570 mT/m/ms and maximum strength 660 mT/m, and the radiofrequency (RF) transmission and reception were achieved with a 72 mm inner-diameter quadrature RF coil. The T2w image for each specimen was acquired with a 3D Turbo RARE sequence: TE = 36 ms, TR = 2000 ms, NEX = 6, matrix size = 240×180×267, resolution = 0.3×0.3×0.3 mm, FOV = 72.0×54.0×80.1 mm. The multi-shell dMRI was acquired with a 3D multi-segment echo planar imaging diffusion weighted sequence: TE = 23 ms, TR = 200 ms, bandwidth = 178.5 kHz, NEX = 7, matrix size = 148×120×160, resolution = 0.45×0.45×0.45 mm, FOV = 66.6×54.0×72.0 mm. For 7 of the 8 monkeys, a total of 186 diffusion-weighted images (DWI) with 2 shells were collected, including 6 non-diffusion images for b = 0 s/mm^2^, 60 directions for b = 2400 s/mm^2^ and 120 directions for b = 4800 s/mm^2^, with a total scan duration of about 243 hours. The scanning for the last monkey consisted of 6 b = 0 s/mm^2^, 30 b = 2400 s/mm^2^, and 90 b = 4800 s/mm^2^ DWI data and lasted about 145 hours.

The <u>Oxford-I</u> and <u>Oxford-II</u> datasets were both from an open resource, PRIME-DE^3, 38, 40^. The Oxford-I dataset included 4 male and 2 female ex vivo subjects. Briefly, it consisted of 16 non-diffusion and 128 b = 4000 s/mm^2^ volumes with 0.6 mm isotropic resolution. The Oxford-II dataset consisted of 20 male in vivo macaque monkeys with 0.5 mm isotropic T1w and 2 mm isotropic resting-state images (TR = 2 s, 1600 volumes). The in vivo TVB dataset was obtained from another public resource^39^ that included 8 male macaque monkeys and consisted of 0.5 mm isotropic T1w images and 1.0 mm isotropic DWI data with 64 directions, b = 1000 s/mm^2^.

#### MRI, Tracer Injection, and Histology of R04

Prior to the surgery, the macaque brain (ID: R04, female, 11 years old, 6.75 kg) was scanned on a 3.0T Simens Prisma scanner to acquire in vivo T1w and T2w images. The T1w images were acquired with a magnetization prepared rapid gradient-echo (MPRAGE) pulse sequence: TE = 2.8 ms, TR = 2.2 s, matrix size = 240×300×320, resolution = 0.5×0.5×0.5 mm, FOV = 120.0×150.0×160.0 mm, and the T2w images were acquired with a single slab three-dimensional turbo spin echo (SPACE) sequence: TE = 0.39 s, TR = 3.2 s, matrix size = 240×300×320, resolution = 0.5×0.5×0.5 mm, FOV = 120.0×150.0×160.0 mm. After the establishment of anesthesia, the scalp was incised under the guidance of a coordinate navigation system (Brainsight Vet, Rogue Research, United States) for the subsequent tracer injection. Three retrograde tracers (1% CTB488, CTB555, CTB647, Invitrogen, United States) were injected with the site coordinates pre-determined in structural MRI images under the path planning of the navigation system. Injections were carried out separately through glass pipettes attached to pumps (Nanoliter 2010 Injector, World Precision Instruments, United States) and a controller (Mico4 Micro syringe Pump Controller, World Precision Instruments, United States) with a volume of 500 nl. 15 days after the surgery, the animal was sacrificed and the brain was extracted and fixed following the protocol mentioned above. The brain was then scanned on a 9.4T scanner to acquire a whole-brain dMRI dataset with the same acquisition parameters as the Brainnetome-8 dataset except that the isotropic resolution was 0.6 mm^3^ with 4 diffusion gradient directions b = 0 s/mm^2^ and 60 b = 4800 s/mm^2^. After the cerebellum was removed from the brain tissue, sucrose protection was utilized, and the tissue was frozen with dry ice and embedded in optimal cutting temperature compound (OCT) before sectioning. Coronal sections with a thickness of 40 μm were obtained using a cryotome (Cryostar NX70, Thermo Fisher, United States) and block-face images were acquired for all sections, using a camera. One in every 12 consecutive sections was selected for Nissl staining to visualize the cell bodies, following the standard procedure. One in every 6 sections was used for fluorescence imaging. Each selected section was imaged with the microscopy (Axio Scan. Z1, Zeiss, Germany) at 0.22 μm in-plane resolution. For those sections with severe artifacts, whenever possible the neighboring sections were chosen for imaging as alternatives.

### Preprocessing

For the ex vivo subjects, the T2w or b0 images were contrast inverted^33^ as a fake T1w to reconstruct the surfaces via the HCP-NHP pipelines^87^; for the in vivo subjects, the surfaces were reconstructed using the T1w structural images directly. Then those surface data were all resampled to 10k and 32k for the subsequent analysis. Additionally, the high-contrast b0 image of a post-mortem template named CIVM^88^ was utilized to reconstruct the CIVM surface for the subsequent definition of seed regions (Fig. 3A).

Preprocessing of the diffusion MRI data was performed in the same way across all the subjects. Specifically, eddy current correction, unringing, and denoising were performed using FSL^89^ and MRtrix3^90^. Finally, fiber orientation distributions (maximal number of fibers per voxel = 3) were estimated using Bedpostx in FSL. The uncertainty provided by Bedpostx characterized the width of the distribution for each orientation. Specially, the averaged b0 images underwent intensity bias correction using N4BiasCorrect^91^ for registration to the CIVM template.

A recently available fMRI pipeline^19^ was adapted to preprocess the resting-state fMRI data following the preprocessing process in published studies^92, 93^. Briefly, it included despiking, motion correction, global scaling, and nuisance regression of the white matter (WM) and cerebrospinal fluid (CSF) signals, bandpass filtering (0.01–0.1 Hz), and co-registration to native anatomical space. Then the denoised resting-state fMRI data was projected to the CIVM midthickness surface and smoothed with FHWM = 3 mm.

The restacked 3D block-face image was used as an intermediate for the coregistration between the dMRI and section images. The Macaque Brainnetome Atlas was first non-linearly mapped to the R04 MRI space and then linearly aligned to block-face space using the ANTs algorithm. A coarse linear alignment between the corresponding section and the block-face image roughly located the brain regions of the MacBNA, and the parcellations were then manually inspected and corrected.

### Connectivity-based Parcellation

#### Initial Seed Region Definition

Following the protocols that we used in constructing the Human Brainnetome Atlas^26^, we identified the initial parcellation units of the macaque brain using previous macaque atlases^16, 18, 88, 94^ or delineations of the macaque brain^95^. Splitting the cortical regions was based on landmarks, such as large sulci and gyri, accompanied by some cytoarchitecturally defined boundaries as a complement (Supplement Fig. S1, Tab. S1). The initial seed regions of the subcortex were solely determined by referring to Paxinos et al.’s^18^ rhesus monkey brain atlas. All the regions were delineated in the CIVM template, and then the cortical ones were projected to the CIVM mid-surface to obtain a surface-based segmentation of 21 regions (Fig. 3A).

#### Surface-based Probabilistic Tractography and Connectivity-Based Parcellation

With 21 cortical seed regions delineated on the surface, a surface-based probabilistic tractography was performed using probtrackx2 in FSL. Taking an initial region as a seed mask (Fig. 3B), 10000 streamlines originating from each vertex were sampled, with a 0.2 curvature constraint, 3200 steps per sample, and a 0.5 mm step length to estimate the vertex-wise connectivity profile. The whole-brain connectivity was smoothed with the tracking target mask resampled to 2 mm to reduce the noise (Fig. 3C). A similarity matrix between each seed vertex was derived from the connectivity matrix and then group-wise averaged. Spectral clustering was used to define the distinct subregions (Fig. 3D) as in our previous study of the human brain^26^. Subcortical regions were parcellated in a similar way, but using tracking in volume space.

#### Strategies for Choosing Optimal Cluster Number and Hierarchical Parcellation

To obtain a reliable parcellation of each region, we selected the local maxima as the optimal cluster number using a weighted score obtained by aggregating three characteristic cluster indices, Cramér’s V (CV), topological distance (Tpd), and hierarchy index (Hi). To validate the robustness of the optimal cluster number, a leave-one-out strategy was used to replicate the identification of local maxima with one subject excluded each time, and we then counted the frequency of each cluster number as a local maximum. These cluster indices were described in detail in Li et al.’s study^96^. An entropy-based aggregate method^97^ was utilized to combine a pairwise CV between individuals, a group-level Tpd between hemispheres, and a group-level Hi of each hemisphere into one. The aggregate score of a cluster number *i* was computed according to:

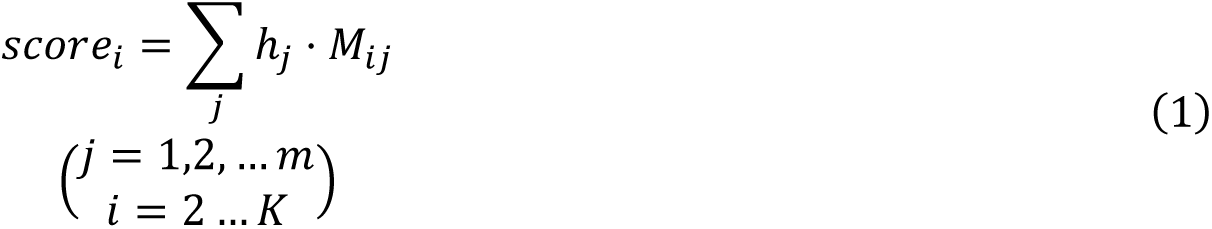

where *Mij* is the value of cluster *i* of the *j^th^*index, and *h_j_* is the weight of the *j^th^* index. Here, *m* is equal to 3 because CV, Tpd, and Hi were all three used, and *K* is the maximal cluster number. The weight *h_j_* was computed as:

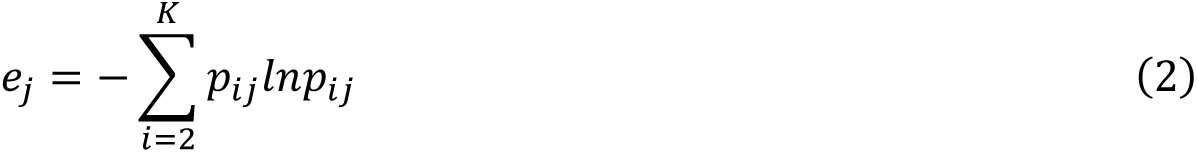

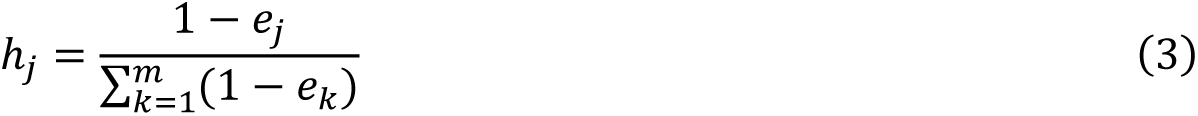

where *p_ij_ is* the normalized *M_ij_* to ensure 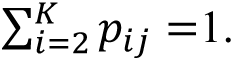

A hierarchical parcellation strategy was employed on three initial ROIs (i.e., IPL, STG, and CG). For example, the IPL was parcellated into a sulcal part with four subregions and a gyral part with only one subregion (Fig. 3F), to which the strategy of parcellation was applied again to obtain a finer pattern.

#### Mapping to Volume and NMT Template

Because the parcellation of the cortex was defined on the brain surface, mapping to the volume space was needed to provide a more general application. Based on the characteristic of columnar anisotropy in the neocortex^27^, each voxel was assigned to the label of its closest vertex along the direction perpendicular to the cortical surface. A manual correction and smoothing operation were applied to fix some errors, such as discontinuities or outliers of the parcellation. Since the new atlas was constructed on an ex vivo CIVM template, we also mapped it to the well-known NMT in vivo template^98^ using a nonlinear registration provided by ANTs^99^ so that our atlas could be used in either setting.

### Structural/Functional Connections and Gradients

For each voxel within a brain region of the MacBNA, a probabilistic tractography using 10000 samples was performed as mentioned above to obtain its structural connectivity to the whole brain. Then, we obtained the structural connectivity of a region with the whole brain by averaging the structural connectivity across all the voxels within it. A group-wise structural connectivity map was calculated by normalizing the connectivity of each subject to standard space and then averaging across all the subjects. For the structural connectome, we counted the connected streamlines as the weight of the edge between regions of the MacBNA for each subject. Then, the connectome was normalized and averaged across the subjects.

The functional connectivity of each region with the whole brain was measured by the Pearson’s correlation coefficient (Fisher’s z-transformed) between its averaged preprocessed time series and that of each voxel of the whole brain. The average group-wise functional connectivity across subjects was tested for statistical significance with a one-sample *t* test (FDR corrected at *P* < .01). For the functional connectome, the parcel-wise Pearson’s correlation was calculated as the functional connection between regions and was then Fisher’s *z*-transformed. A one-sample *t* test was used to determine the statistical significance on the group level (FDR corrected at *P* < .01).

Diffusion map^46^ was used on the structural and functional connectivity to obtain gradients at the vertex wise level. The gradients of the seed region were derived from the similarity matrix between the connectivity of each voxel within the brain region to explore the pattern of changes along specific directions as a complement to and explanation of the parcellation. We selected the top 20 components in the embedding algorithm and visualized the gradients with t-SNE^47^ in a 2D dimension.

### Evaluation with Tracer Data

Tracer-derived connections are the standard for validating connections identified by diffusion MRI. We evaluated the MacBNA parcellation and the dMRI connections using the collected tracer data from R04 and an open-access tracer resource^34, 43^.

First, we compared the retrograde tracers of the three injected brain regions from the MacBNA with the corresponding connections derived from the dMRI data from the same animal (R04). The seed ROIs were identified as the intersection of a sphere (radius = 3 mm) centered at the coordinates of each injection site and the white/gray surface. 10000 probabilistic streamlines were sampled from each voxel within each seed ROI to estimate the structural connections, as mentioned above, using the dMRI data from R04. The group-wise structural connections of the Brainnetome-8 were also compared with the tracer data from R04.

Furthermore, an open tracer resource of macaque data^34, 43^ was used to evaluate the quality of the Brainnetome-8 dMRI data and to validate the structural connectome derived from MacBNA. The original 29×91 tracer connections and the parcellation with 91 cortical regions were used to calculate the correlation with the structural connections using the Brainnetome-8, Oxford-I, and TVB, taking distances between regions into consideration. The distances between the regions of the parcellation from Markov et al.^43^ were released in a previous study^100^, which identified the minimum white matter trajectories. Furthermore, we mapped the injection sites to the CIVM template surface to determine the target regions in our atlas. Nine target regions from Markov et al.^43^ were excluded because the related injection sites did not satisfy a one-to-one mapping, i.e., multiple sites projected to the same region of our atlas or were located near the boundary. The tracer connectome was mapped to our atlas simply based on the overlap ratio with Markov et al.’s^43^ parcellation. This was possible because Donahue et al.^34^ provided the tracer coordinates that Markov et al.^43^ had used in their study, enabling us to validate our atlas with Markov et al.’s tracer data. Thus, we obtained a 20×124 transformed tracer-derived connection matrix to compare with the structural connections from MacBNA while considering the areal distances (Fig. S15). Finally, a 20×20 subgraph that represented the connections between the injected regions was extracted from the adapted tracer matrix to calculate the correlation with the dMRI ones. The distances between the regions of MacBNA were determined as the mean distance of the streamlines using dMRI tractography. The dMRI connection matrix was averaged across hemispheres and fractionally scaled and log normalized for consistency with the tracer connections^34^. Both the tracer and dMRI connection matrices were symmetrized before the correlation to keep them comparable with each other. The connection matrices were also binarized for the calculation of the area-under-curve score to evaluate the sensitivity and specificity of the dMRI connections.

### Comparisons with Existing Parcellations

Five digital parcellations based on cytoarchitecture were used to compare with MacBNA. The alternative parcellations include: (1) Paxinos et al.’s brain atlas of macaques, which has a digital 3D version^88^ (PTH09); (2) The fifth-level (CHARM_5) and (3) the sixth-level parcellation (CARHM_6) of a hierarchical atlas^19^; (4) Saleem et al.’s digital atlas based on five different histological stains^44^ (Saleem); (5) The parcellation that was used in Markov et al.’s study^43^ to navigate the injection sites of tracer for which we used its digital version^34^ (Markov).

We compared the six parcellations using a homogeneous metric, DCBC. The basic idea underlying DCBC^101^ is that any two vertices/voxels within the same region should have more similarity between their structural connectivity profiles than those belonging to distinct regions. Taking the spatial smoothness of brain data into consideration, DCBC binned each vertex/voxel pair (bin step = 1 mm) based on their geometric distance ranging from 0 to 100 mm and then compared the similarity between the within and between-parcel within each bin. A higher DCBC indicates that the parcellation reflected a more homogeneous topography based on structural connectivity. The DCBC of MacBNA was compared with that of the other parcellations across subjects using paired *t* tests and was then FDR corrected to test the statistical significance.

### Cytoarchitectonic Analysis

#### Cytoarchitectonic Boundary

The digitized images of the Nissl-stained sections were used for the cytoarchitectonic analysis as Amunts et al.^10^ did. Briefly, the images were down-sampled to 22 μm and smoothed to reduce noise and artifacts. The optical density profiles of the regions of interest, which captured the laminar changes, were extracted orthogonally to the surface. The features vectors extracted from the profiles were used to measure the Mahalanobis distance between neighboring blocks. A sliding window procedure (11 <= block size <= 24) determined the peak (maxima) of the distance function. A Hotelling’s T^2^ test was used to confirm the statistical significance of the peak. The peak locations identified using multiple block sizes were also evaluated by comparing them with maxima locations at neighboring stained sections to exclude the false positive peaks caused by artifacts or outliers (Figs. S3, S6).

#### Glial Framework

The glial framework was obtained by orientation estimation and fiber tractography using Schurr and Mezer’s^37^ method. In short, a structure tensor analysis was used to fit the local fiber orientations according to the spatial distribution of the glial cells. The tile size was set at 44 microns (i.e., 200 pixels) with ρ = 15 and σ = 3 as the default. A white matter mask was generated using a semi-automatic annotation tool^102^ based on the contrast between the gray and white matter in each Nissl slice. The principal direction estimated using dtifit in FSL with dMRI data from the same monkey was aligned to block-face space for comparison with the orientation obtained from the glial organization. The white matter pathway was reconstructed using fiber assigned by continuous tracking (FACT)^103^, a deterministic tractography algorithm in MRtirx3^90^. Step size was set at 3 voxels and the maximal angle was 60°. The seed mask was located in the white matter and 25 streamlines were seeded for each voxel.

### Cross-species Structural Divergence Based on Brainnetome Atlases and Its Association with Transcriptome

42 homologous tracts (38 association and 4 commissure) were carefully dissected automatically with XTRACT^104^ to characterize the connectivity blueprint^48, 104^ of each region for each macaque in the Brainnetome-8 and each human among 500 right-handed subjects randomly selected from the HCP-YA dataset^105^ with minimal preprocessing^106^. Each vertex of the white/gray matter surface was seeded for probabilistic tractography with the whole brain as the target to generate a vertex-wise connectivity matrix. Then, it was multiplied by the vectorized tract map that described the trajectory of each tract in the brain and was normalized to get the connectivity profiles of each vertex with the 42 tracts. Finally, we averaged the connectivity profiles of the vertices within each region of the MacBNA and HumBNA to form the regional connectivity blueprints across subjects. In other words, the connectivity blueprints represent the connection of each cortical region to the 42 major white bundles. The parameters were set as a default of the XTRACT tool. The symmetric Kullback Leibler (KL) divergence was used to measure the similarity of the group-wise connectivity blueprints between each region of MacBNA and each region of HumBNA. The divergence map represents the minimal KL divergence of each region in the human from the macaque regions. The divergence map was compared using the Spearman’s correlation with another brain maps that are provided in neuromaps^49^.

The regional gene expression of the HumBNA was estimated with the Allen Human Brain Atlas (AHBH) transcriptome dataset^50^ using abagen^107^, resulting in 15633 remaining genes. Only the left hemisphere was considered since the AHBH has probe samples in the right hemisphere for only two subjects^108^. A partial least squares regression (PLSR) between the divergence map and gene expression was performed to characterize the genetic mechanism underlying the structural variation between the two species. To avoid overfitting, only the genes that significantly correlated with the divergence map (*P* < .05) were retained for the subsequent PLSR analysis. PLSR has been widely used to bridge the relationship between neuroimaging and the transcriptome because it determines the most predictive genes with a weighted score of the neuroimaging map^109–111^. 10000 surrogate maps, which were generated by Brainsmash^112^, were used to ensure that the explained variance of the first PLSR component (PLS1 score) was significantly higher than that generated by chance. The correlation between the divergence map and the PLS1 score was examined as described above. Then, 1000 bootstrapping was used to estimate the error of the weight of each gene, and the normalized weight (Z) of each gene was denoted as the weight divided by the estimated error^111^. We quantified the enrichment of the statistically significant genes (|Z| > 3, 2253 genes remained) in the disease-related gene sets, which we obtained from DisGeNet^113^, with all the human genes as the background using Metascape^114^ software. Finally, we tested whether these significant genes overlapped more with human-accelerated genes (HAR genes)^52^ using Fisher’s exact test.

### Statistical Significance using Permutated Brain Data

Due to the spatial autocorrelation of the brain data, we tested the statistical significance (*P*_spin_) of the findings by generating surrogate brain maps that preserved the spatial autocorrelation using BrainSMASH^112^. The number of permutation samples was 10000, if not specifically stated.

## Abbreviation

IPL: inferior parietal lobe
IPL.gr: inferior parietal lobule, gyral rostral part inferior
IPL.sr: parietal lobe, sulcal rostral part
FPO.ci: frontal parietal operculum, caudointermediate part
PM.rd: premotor cortex, rostrodorsal part
PM.cd: premotor cortex, caudodorsal part
PM.i: premotor cortex, intermediate part
PM.cv: premotor cortex, caudoventral part
PM.rv: premotor cortex, rostroventral part
PM.rm: premotor cortex, rostromedial part
PM.cm: premotor cortex, caudomedial part
SFG.ri: superior frontal gyrus, rostrointermediate part
IFG.cd: inferior frontal gyrus, caudodorsal part inferior
IFG.cv: frontal gyrus, caudoventral part
OrG: orbital gyrus
FP: frontal pole
FP.d: frontal pole, dorsal part
M1: primary motor cortex

## Data and Code Availability

The ex vivo MRI data of the Brainnetom-8, the MRI images and sections of another postmortem monkey, and the code of the parcellation pipeline based on the surface will be openly available at https://molicaca.github.io/atlas/mcqatlas.html upon acceptance.

## Acknowledgements

This work was partially supported by the Science and Technology Innovation 2030 - Brain Science and Brain Inspired Intelligence Project (Grant No. 2021ZD0200200), Natural Science Foundation of China (Grant Nos. 82151307, 82202253, and 31620103905), Strategic Priority Research Program of the Chinese Academy of Sciences (XDB32030207), and Science Frontier Program of the Chinese Academy of Sciences (grant No. QYZDJ-SSW-SMC019).

